# T-scope V4: miniaturized microscope for optogenetic tagging in freely behaving animals

**DOI:** 10.1101/2024.10.07.616920

**Authors:** Yuteng Wang, Pablo Vergara, Satoshi Hasegawa, Naoki Tomita, Yoan Cherasse, Toshie Naoi, Takeshi Sakurai, Yuki Sugaya, Masanobu Kano, Masanori Sakaguchi

## Abstract

A miniaturized microscope (i.e., miniscope) enables the imaging of neuronal activity using calcium sensors while simultaneously manipulating that activity using opsins in freely moving animals. However, many miniscopes use light-emitting diodes with broadband emission, leading to unintentional opsin stimulation by light intended solely for calcium sensor activation (a phenomenon referred to as “biological crosstalk”). To address this issue, we previously developed a miniscope including a port for chosen light sources, such as lasers, by restructuring the open-source UCLA Miniscope v3. However, targeting the same neuronal soma for both excitable opsin stimulation and calcium sensor imaging remained a challenge. Here, we integrated features from the UCLA Miniscope v4 into our new T-scope V4 miniscope. In optogenetic tagging experiments, we demonstrated that a 445-nm blue laser can be used to image neuronal activity with the calcium sensor GCaMP6s without inadvertently stimulating the ChrimsonR opsin, allowing for simultaneous neuronal activity imaging and manipulation in freely moving mice. Thus, the T-scope V4 can serve as a powerful tool for probing causal relationships between neuronal activity and its function in living animals.

**Highlights:**

- We developed the T-scope V4 that integrates features of the UCLA Miniscope v4
- This miniscope prevents “biological crosstalk” between sensors and opsins
- This tool can help probe causal brain-behavior relationships in living animals

## Introduction

Genetically encoded calcium indicators, such as GCaMP6, offer a robust method for monitoring neural activity by observing changes in fluorescence that correspond to alterations in calcium concentration^1^. Further, calcium imaging combined with optogenetics allows the investigation of neuronal circuit dynamics through the manipulation of neuronal activity^2^. Combining this approach with two-photon microscopy can even enable the manipulation of specific neurons^3^. However, the application of these technologies has been limited due to the need for an elaborate setup and/or immobilization of the animal’s head, especially in freely behaving settings^4,5^.

Recently, simultaneous calcium imaging and optogenetic manipulation in freely behaving animals has become possible using a one-photon miniaturized microscope^6^. These miniscopes often employ light-emitting diodes (LEDs) with broadband emission, leading to unintentional opsin stimulation by light intended solely for calcium sensor activation (i.e., “biological crosstalk”)^6^. Although the recent development of red-shifted channelrhodopsins show promise for alleviating this issue, their spectral properties do not eliminate the biological crosstalk occurring under persistent calcium imaging using blue light^7,8^. This unresolved issue can potentially introduce confounding variables. For instance, when using the combination of ChrimsonR (a red-shifted light-gated ion channel) and GCaMP6, lower power blue LED light can trigger depolarization in ChrimsonR-expressing neurons if the light is directly delivered to their soma^1,6^.

To address this issue, we previously modified the UCLA Miniscope v3 to accept flexible light sources, including lasers that emit monochromatic light for concurrent imaging and optogenetic silencing^9^. However, evidence demonstrating its capability for simultaneous imaging and excitation targeting the same neuronal soma was lacking. Here, we created the T-scope V4 miniscope based on open-source UCLA Miniscope v4 architecture^10^ to overcome biological crosstalk. Using an optogenetic tagging method, we show that a 445-nm blue laser markedly diminished unintentional ChrimsonR stimulation. Therefore, the T-scope V4 offers the opportunity to examine the same neuronal population over extended periods, which can allow investigation of their causal relationship with behavior in freely moving animals.

## Materials and Methods

### Animals

All animal experiments were approved by the University of Tsukuba Institutional Animal Care and Use Committee. Mice were maintained in insulated chambers at an ambient temperature of 23.5 ± 2.0°C under a 12-h light/dark cycle with ad libitum access to food and water according to institutional guidelines. Mice (Jackson Laboratory, Sacramento, CA, USA) harboring TIGRE-Ins-TRE-loxP-stop-loxP(LSL)-GCaMP6s (Ai94D, stock #024104) were backcrossed in a C57BL6/J background more than 5 times.

### Virus preparation

The following virus vectors were used for experiments: adeno-associated virus (AAV)1-Syn-Flex-ChrimsonR-Tdtomato (Addgene #62723), AAV2retro-cFos-tTA-pA (Addgene #66794), and AAV2retro-CaMKII-0.4-Cre (Addgene #105558). AAV vectors were prepared as previously described^11^.

### Virus injection and lens implantation

Surgery was performed at 9-12 weeks of age. Mice were anesthetized with isoflurane and fixed in a stereotaxic frame (Stoelting). The height of bregma and lambda were adjusted to be equal. For granular neuron labeling, 70 nl of a 1:1 volume mixture of AAV solution was injected into the dorsal hippocampus at anterior-posterior (AP) - 2.0 mm, medial-lateral (ML) +1.2 mm, and dorsal-ventral (DV) -1.7 mm relative to bregma. A female mouse was used to induce ChrimsonR and GCaMP6s expression, while a male mouse was used to induce GCaMP6s. Mice were allowed to recover for 1 week before lens implantation.

One week after virus injection, microendoscope lens (1-mm diameter, 4-mm length, Inscopix) implantation and baseplate installation were performed as previously described^12^. Briefly, the lens was placed at AP -2.0 mm and ML +1.25 mm relative to bregma and a depth of 1.53 mm below the dura. One week after lens placement, the baseplate for the miniscope was attached above the implanted microendoscope lens. After baseplate surgery, mice were habituated to the attached dummy miniscope for at least 7 days before recording.

### T-scope V4 construction

As previously described, we modified components related to the light path based on the original construction of the UCLA open-source V4 Miniscope. We added a port connected to the excitation module for laser delivery instead of the original LED source. This port included a sputtered enhanced silver reflective mirror (21015, Chroma) and light-shaping diffusers (LSD20PC10-12, Luminit). To allow the combination of GCaMP imaging and ChrimsonR modulation, we replaced the dichroic mirror (ZT530/55dcbp, Chroma) in the emission module. These modifications were made to block out any unforeseen light to reduce crosstalk and background noise. The 3D structural data is available in Mendeley Data (DOI: 10.17632/5g8xf7jgwm.1) (temporary link for review purposes: https://data.mendeley.com/v1/datasets/5g8xf7jgwm/draft#folder-9c76ec4d-09f8-4053-94d4-1f0aa3b229de).

### Calcium imaging and manipulation

For the optogenetic tagging experiment, a blue laser (445 nm, custom-made, 0.5-1.1 mW at the bottom of the object lens) was used for calcium imaging, and an orange laser (589 nm, Shanghai Lasers, China, 0.5 mW at the bottom of the object lens) was set to emit light pulses at 10 Hz (50 ms on/off cycles) for ChrimsonR stimulation. The laser light was combined and sent through a single optic patch cable (Thorlabs).

The optogenetic tagging experiment was performed for a duration of 300 s. Throughout the entire session, a blue laser was used to continuously record changes in calcium levels. In parallel, an orange laser emitted a 1-s pulse at the conclusion of each 30-s imaging block, resulting in a total of 10 pulses during the 300-s session.

### Brain slice characterization

After completion of the experiment, mice were perfused transcardially with phosphate-buffered saline (0.1 M) and 4% paraformaldehyde. Brains were removed, fixed overnight in paraformaldehyde, and transferred to phosphate-buffered saline. Coronal sections (30 μm) were cut using a vibratome (VT1200S, Leica). Sections were mounted on slides with mounting medium containing DAPI (Merck). After fixation, imaging analysis was performed from sequential z-series scans with a Leica TCS SP8+ confocal microscope.

### Calcium imaging data analysis

Calcium imaging recordings were processed using the CaliAli pipeline^13^ (https://github.com/CaliAli-PV/CaliAli). Briefly, raw calcium imaging videos were spatially downsampled by a factor of 2. Motion correction was then applied using translation and log-demon registration to correct non-rigid deformations.

Displacement fields were estimated from a filtered version of the video in which blood vessels and neurons were enhanced using hessian-filters and anisotropic diffusion, respectively. These estimated displacement fields were subsequently applied to the downsampled video. The motion-corrected video was then detrended and neural signals were enhanced using the neural enhancing module described in MIN1PIPE^14^. Neuron tracking across sessions was performed by aligning and concatenating the videos, utilizing blood vessels and neurons as reference points to correct for inter-session misalignment. Finally, the concatenated video was processed using the CNMF-E^15^ implementation described in CaliAli.

### Statistical analysis

Statistical analysis was performed using GraphPad Prism version 7.04 for Windows (GraphPad Software) and MATLAB (Mathworks). Error bars represent the 95% confidence interval of the mean. Shaded-in error bars were derived using a bias-corrected and accelerated bootstrap methodology. Type I error was set at 0.05.

## Results

We modified components related to the light input of the UCLA open-source Miniscope V4 architecture to allow any laser light source to replace the LED (Fig. 1a). We added an optic ferrule port to the excitation module to facilitate delivery of the light source. This module incorporates a reflective mirror and light-shaping diffusers. We also replaced the dichroic mirror for optimization of simultaneous calcium imaging and optogenetic stimulation. Although these modifications increased the overall weight of the miniscope to ∼4 g, no noticeable changes were observed in mouse behavior following a 1-week habituation period with a dummy miniscope.

**Figure 1.**
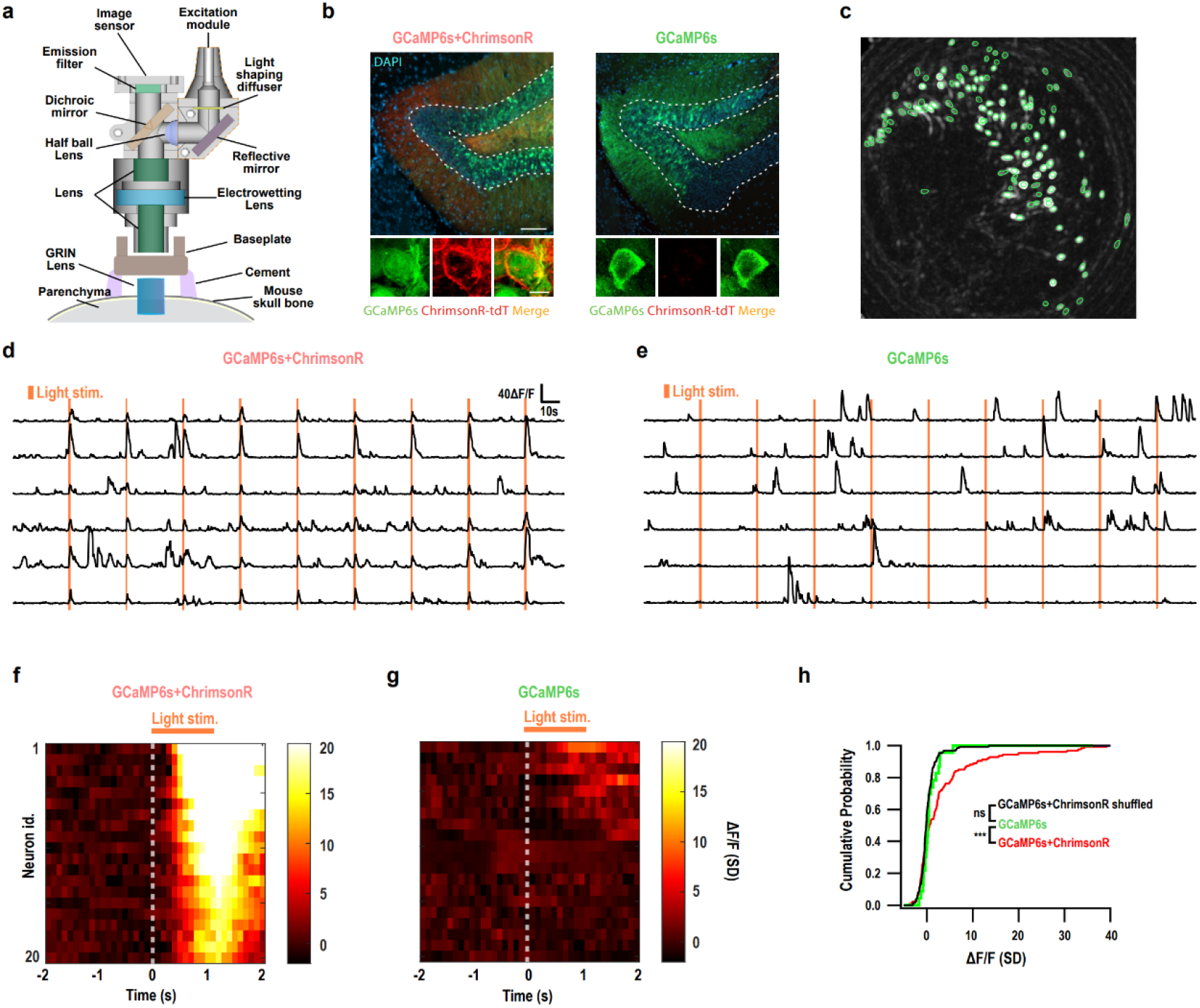
Optogenetic tagging experiment using a laser-based miniscope. **a**, Cross-sectional drawing of the T-scope V4. **b**, Representative images of GCaMP6s and ChrimsonR expression in granular neurons (DAPI, blue; ChrimsonR-tdTomato, red; GCaMP6s, green; scale bars: top, 100 μm; bottom, 5 μm). **c**, Representative neuron contours. **d**,**e**, Representative calcium traces obtained during 589-nm light stimulation in GCaMP6s- and GCaMp6s+ChrimsonR-expressing mice. **f**,**g**, Heatmaps of standardized peristimulus responses for the 20 neurons with the largest reactions during light stimulation in GCaMP6s- and GCaMp6s+ChrimsonR-expressing mice. **h**, Cumulative probability plot depicting the distribution of average changes in *ΔF/F* during light stimulation in GCaMP6s- and GCaMp6s+ChrimsonR-expressing mice, with shuffled data for GCaMP6s+ChrimsonR (random circular shifts of calcium traces; Kolmogorov-Smirnov test with Bonferroni correction. GC (n = 23) vs. GC-Chr (n = 126), P = 0.0010; GC (n = 23) vs GC-Chr Shuffled (n = 126), P = 0.18).

To validate the simultaneous imaging and optogenetic manipulation capabilities of our T-scope V4, we performed an optogenetic tagging experiment in the hippocampal dentate gyrus. We used one AAV to induce expression of GCaMP6s to monitor neural activity^1^ and another AAV to induce expression of ChrimsonR to manipulate neural activity using orange light stimulation in a mouse^7^ (Fig. 1b). A control mouse expressed only GCaMP6s. A GRIN lens was placed above the dentate gyrus to deliver light and correct emission from the calcium sensor (Fig. 1c). To mitigate the risk of unintentional ChrimsonR stimulation during calcium imaging, we used a 445-nm blue laser to stimulate GCaMP6s (Fig. 1b). The optogenetic tagging experiment comprised 10 trials of imaging and stimulation, with each trial consisting of a 29-s baseline recording followed by 1-s stimulation using a 589-nm orange laser. In the mouse co-expressing GCaMP6s and ChrimsonR, a subset of neurons exhibited prominent calcium events in response to each orange laser stimulation (Fig. 1d, f).

This optical response upon orange laser stimulation was not observed in the control mouse (Fig. 1e, g). Moreover, the distribution of average fluorescence response in the mouse co-expressing GCaMP6s and ChrimsonR was significantly different from a null distribution obtained by random circular shift of calcium traces, whereas this null distribution did not differ significantly from the distribution in the control mouse (Fig. 1h). These results demonstrate the successful optogenetic tagging of neurons with the combination of GCaMP6s and ChrimsonR using T-scope V4.

## Discussion

The major limitation of combining fluorescent calcium sensors with excitatory opsins is spectral overlap. This arises due to the generally broad range of sensitive wavelengths of opsins, leading to possible unintentional opsin activation during calcium imaging. Although reducing the intensity of blue light may lower the probability of unintentional opsin actuation, it can also reduce the peak-to-noise ratio of the signal from calcium sensors. This, in turn, can pose difficulties in extracting individual calcium events. In contrast to LED light sources commonly utilized in miniscopes, single-wavelength lasers are potential alternatives that could mitigate this issue. In this study, we present a miniscope capable of accepting combinations of light sources, including lasers, for simultaneous calcium imaging and optogenetics in free-moving mice. The flexibility of the light source also allows the selection of other combinations of fluorescent calcium indicators and optogenetic tools. By using the combination of GCaMP6s and ChrimsonR, we successfully induced neuronal activity and simultaneously detected corresponding calcium signals. This tool could enable the tracking of the activity of genetically defined neuronal populations to determine their functional causal links to behavior in unconstrained animals in future studies.

## Abbreviations

AAV: adeno-associated virus
LED: light-emitting diode

## Data availability

Source data is available at Mendeley Data (DOI: 10.17632/5g8xf7jgwm.1) (temporary link for review purposes: https://data.mendeley.com/v1/datasets/5g8xf7jgwm/draft#folder-9c76ec4d-09f8-4053-94d4-1f0aa3b229de).

## Acknowledgements

We thank M. Sakurai for secretarial support and K. Akers for English proofing. We also thank all WPI-IIIS members.

